# Modeling Retinitis Pigmentosa 59: *Dhdds* T206A and *Dhdds* K42E knock-in mutant mice are phenotypically similar

**DOI:** 10.1101/2025.01.13.632845

**Authors:** Mai N. Nguyen, Dibyendu Chakraborty, Jeffrey Messinger, David M. Sherry, Steven J. Fliesler, Steven J. Pittler

## Abstract

<u>D</u>e<u>h</u>ydro<u>d</u>olichyl <u>d</u>iphosphate <u>s</u>ynthase (DHDDS) is an essential enzyme required for several forms of protein glycosylation in all eukaryotic cells. Surprisingly, three mutant alleles, (*Dhdds*^K42E/K42E^ (K42E/K42E), *Dhdds*^T206A/K42E^ (T206A/K42E), and found in only one patient, *Dhdds*^R98W/K42E^ (R98W/K42E) have been reported that cause non-syndromic retinitis pigmentosa (RP59), an inherited retinal degeneration (IRD). Because T206A was only observed heterozygously with the K42E allele in RP59 patients, we used CRISPR/CAS9 technology to generate T206A/T206A, and subsequently T206A/K42E alleles in mice to assess the contribution of the T206A allele to the disease phenotype, to model the human disease, and to compare resulting phenotypes to our homozygous K42E mouse model. By postnatal (PN) 12-mo, T206A/K42E mice exhibit significant reduction of inner nuclear layer thickness as was observed in K42E/K42E mice. No change in outer nuclear layer thickness is observed in all mutant phenotypes up to PN 12 mo. Electroretinography (ERG) showed a significantly reduced b-wave without a-wave decrement and by PN 3-mo, ERG c- and d-wave responses were significantly attenuated in all phenotypes. Consistent with a reduction in inner nuclear layer thickness seen by OCT and cell loss observed by histology, bipolar and amacrine cell densities were reduced in all *Dhdds* mutant phenotypes compared to PN 8-12 mo age-matched controls. These results indicate that the *DHDDS* T206A allele causes retinal disease independent of the K42E allele, and that there likely is a common disease mechanism involving RP59-associated *DHDDS* mutations. We conclude that the physiological basis of retinal dysfunction in RP59 involves defective signaling in the inner retina resulting in bipolar/amacrine cell degeneration.

## INTRODUCTION

In the dolichol synthesis pathway, two dehydrodolichyl diphosphate synthase (DHDDS; OMIM #608172, EC: 2.5.1.87) subunits partner with two Nogo-B receptor subunits (NgBR; OMIM #610463) to form the *cis*-prenyltransferase (CPT) enzyme complex (Bar-El et al., 2020; Brasher et al., 2015; Edani et al., 2020; Sato et al., 1999). This enzyme complex catalyzes the sequential, repeated addition of five-carbon (C-5) units (isopentenyl diphosphate, IPP) to farnesyl diphosphate (FPP) to form polyprenyl diphosphates, which ultimately are converted to dolichols (Dol) of varying chain lengths (typically Dol-18 to Dol-20). Recently, the “conventional wisdom” regarding Dol formation has been challenged by findings that implicate the formation of aldehydes (polyprenal and dolichal) as obligate intermediates in the synthesis of Dol (Wilson et al., 2024). Dol is required for protein glycosylation (Brasher et al., 2015; Doucey et al., 1998; Kornfeld and Kornfeld, 1985; Sato et al., 1999; Welti, 2013); its phosphorylated derivative (Dol-P) serves as a lipid-soluble carrier for sugar nucleotides and for the formation of oligosaccharide chains that are utilized for protein glycosylation (Aebi, 2013; Cherepanova N, 2016). Disorders arising from defects in any of the genes encoding the more than 30 enzymes in the dolichol and protein glycosylation pathways are grouped together as a family termed “congenital disorders of glycosylation” (CDGs; OMIM #617082) (Cantagrel and Lefeber, 2011; Eklund and Freeze, 2006). Absence of DHDDS is embryonic lethal (Brandwine T, 2021), and more subtle mutations in the enzyme lead to potentially fatal downstream effects in the glycosylation pathway (Cantagrel and Lefeber, 2011; Hemming, 1992; Sabry et al., 2016) or moderate to severe brain disease including epileptic encephalopathies, myoclonus ataxia, or Intellectual Deficit Disorder (IDD), Parkinson’s Disease-like symptoms, and neurodevelopmental disorders (Galosi et al., 2021; Kim J, 2021; Piccolo et al., 2021). Three mutations in the DHDDS gene (K42E, T206A, and R98W) have been reported (Fliesler et al., 2022; Kimchi et al., 2018; Sabry et al., 2016; Zelinger et al., 2011; Zuchner et al., 2011) that cause a recessive form of retinitis pigmentosa called “RP59” (OMIM #613861). Surprisingly, this hereditary retinal disease is *non-syndromic*, *i.e.,* the associated pathologies are restricted to the retina, without any obvious systemic involvement, despite the fact that DHDDS is required for dolichol synthesis and, in turn, for protein glycosylation in *every cell type and tissue* throughout the body.

Retinitis pigmentosa (RP) comprises a heterogeneous group of individually rare genetic disorders of the retina that cause impaired vision and, ultimately, blindness in about 1:4,000 individuals in developed countries (Ali et al., 2017). The classic form of the disease first disrupts the rod photoreceptors, primarily causing reduced night vision (nyctalopia). Peripheral vision is initially affected, due to the dominance of rod photoreceptors (relative to cones) in the periphery, causing “tunnel vision;” but as the disease progresses with time, central vision (mediated by cone photoreceptors) becomes diminished and eventually can result in total blindness (Fahim, 2017). Mutations in more than 70 genes have been linked to classic RP and over 300 genes are involved in inherited retinal diseases (IRDs) (https://sph.uth.edu/retnet/).

We previously reported on a novel murine knock-in model of RP59 (*Dhdds*^K42E/K42E^) that did not exhibit overt signs of photoreceptor degeneration nor altered glycosylation up to postnatal (PN) 12 months (mo) of age. There was, however, a marked increase in anti-GFAP (glial fibrillary acidic protein) immunoreactivity (Ramachandra Rao et al., 2020), indicative of a reactive Müller cell glial response (“gliosis”) to disturbance in the environment of the retina. In-depth examination revealed reduced inner nuclear layer (INL) thickness, retraction of photoreceptor terminals, and “sprouting” of horizontal and bipolar cell processes into the outer nuclear layer (ONL), which are typical of various forms of photoreceptor degeneration (Nguyen et al., 2023). Physiologically, these mutant mice exhibited a “negative b-wave” ERG (*i.e*., ERG b-wave amplitudes below baseline) waveform. Analysis of isoprenoid lipids showed dolichol species having shorter than normal chain lengths compared to those of wildtype (WT) control mice, with the mutants exhibiting a shift to Dol-17/Dol-18 species compared to Dol-18/Dol-19 as the dominant species in WT mice, similar to results reported for Dol species in RP59 patient bodily fluids (Wen et al., 2013). Surprisingly, although RP59 is classified as a CDG, there was no evidence of compromised global protein *N*-glycosylation in that RP59 mouse model (Ramachandra Rao et al., 2020).

In this study, we characterized another *DHDDS* allele – T206A, only found heterozygously with the K42E allele in RP59 patients (Wen et al., 2013). We created a novel *Dhdds*^T206A/K42E^ (T206A/K42E) compound heterozygous mouse line to model the human disease and a homozygous *Dhdds*^T206A/T206A^ line to assess pathogenicity of the T206A mutation. Our results show that the T206A mutant allele itself is pathogenic and that the T206A and K42E alleles cause similar phenotypes that likely arise from the same disease mechanism.

## RESULTS

### Creation of a T206A DHDDS knock-in mouse line using CRISPR/Cas9 technology

DHDDS is a ubiquitous protein encoded by 333 codons distributed across 8 exons and one additional noncoding 5’-exon (**Fig. 1**). The K42E mutation is encoded in exon 3, R98W is in exon 4, and T206A is in exon 6 (**Fig. 1A**). Functional domains of the DHDDS protein are encoded within exons 3-6, including a cofactor binding site for Mg^2+^ (one Mg^2+^ bound to each subunit) and multiple isopentenyl diphosphate binding sites. In this study, the T206A knock-in mouse line was generated using a modified CRISPR/Cas9-based technology. The T206A knock-in mutation was confirmed in T206A/WT mice by mouse tail DNA sequence analysis of a PCR product amplified from the *Dhdds* locus (*black arrow*, **Fig. 1B**) showing an A-to-G nucleotide change. In addition, a silent C-to-A polymorphism was included in the construct to remove the PAM site sequence required by CRISPR for DNA cleavage (**Fig. 1B**).

**Figure 1.**
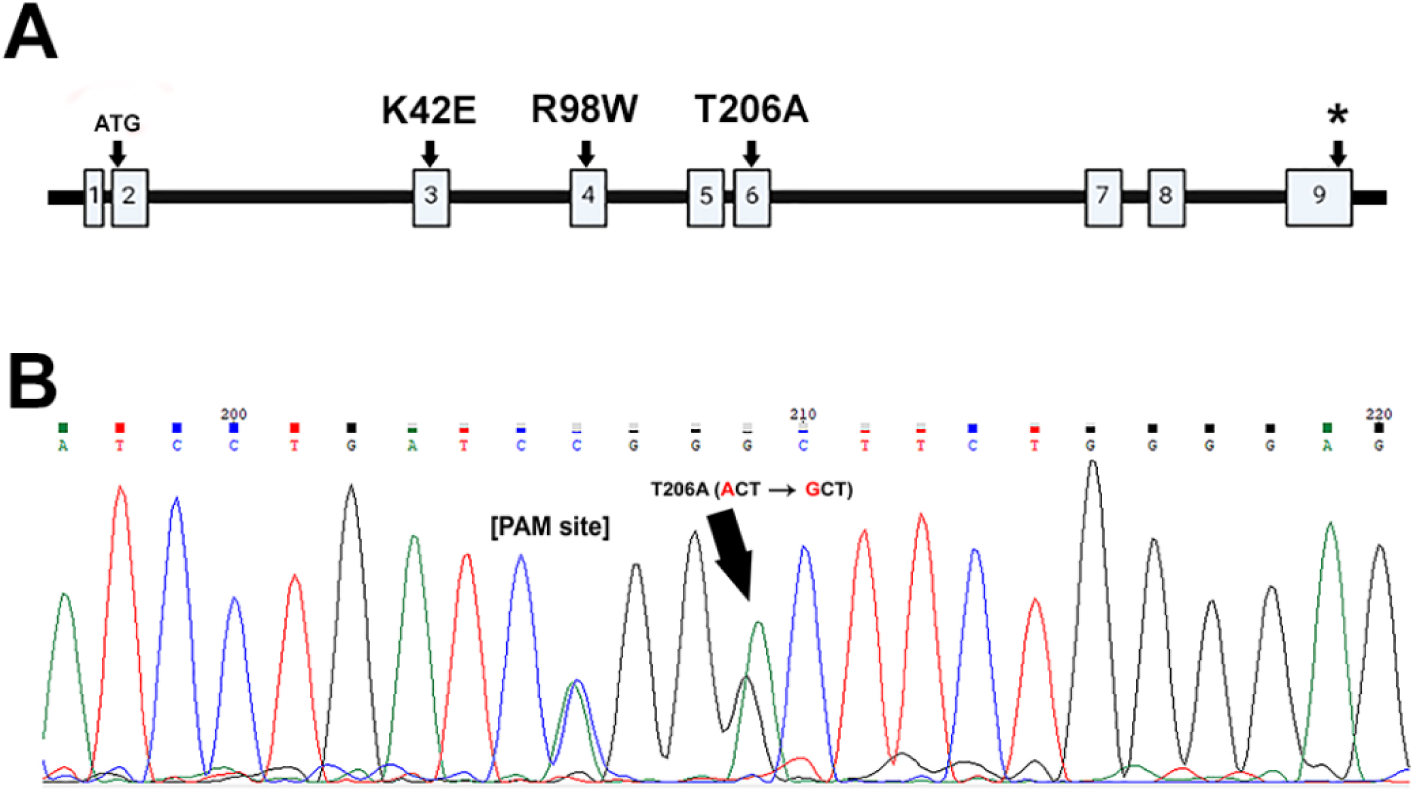
Verification of T206A knock-in mutation. **(A)** A schematic representation of the DHDDS gene, showing its nine exons, with open reading frame ATG start codon encoded in exon 2, K42 in exon 3, R98 in exon 4, T206 in exon 6, and the stop codon (*) in exon 9. Within the 3D structure (not shown), K42 and T206 flank the binding sites, including a single Mg^+^ ion per subunit and multiple isopentenyl units (Bar-El et al., 2020). **(B)** DNA sequence of tail snip DNA from an F0 founder mouse, showing the A-to-G T206A mutation (*black arrow*) and a C-to-A silent heterozygous polymorphism that removes the CRISPR-related PAM recognition site that is required for CRISPR mediated DNA cleavage.

### Retinal morphology of *Dhdds* mutant and WT mice

Transverse sections of epoxy-embedded retinas from PN ≤ 6-mo old mice of genotypes WT, T206A/T206A, T206A/K42E, and K42E/K42E all showed normal retinal layer stratification (**Fig. 2A**). Well-formed outer and inner plexiform layers (OPL and IPL, respectively) were present in all genotypes. Pathologic changes in retinal organization became apparent in all mutant genotypes at older ages (PN ≥ 12-mo; **Fig. 2B**) including ectopic rod cell bodies invading the OPL (displaced PR nuclei, *black arrows,* **Fig. 2B**). Retinal thinning also was evident, particularly in the INL and IPL of T206A/K42E and K42E/K42E mice, indicating loss of inner retinal cells. Other than displaced PR nuclei, inner retinal changes were apparent but less pronounced in T206A/T206A mice compared to the other mutant genotypes containing the K42E allele.

**Figure 2.**
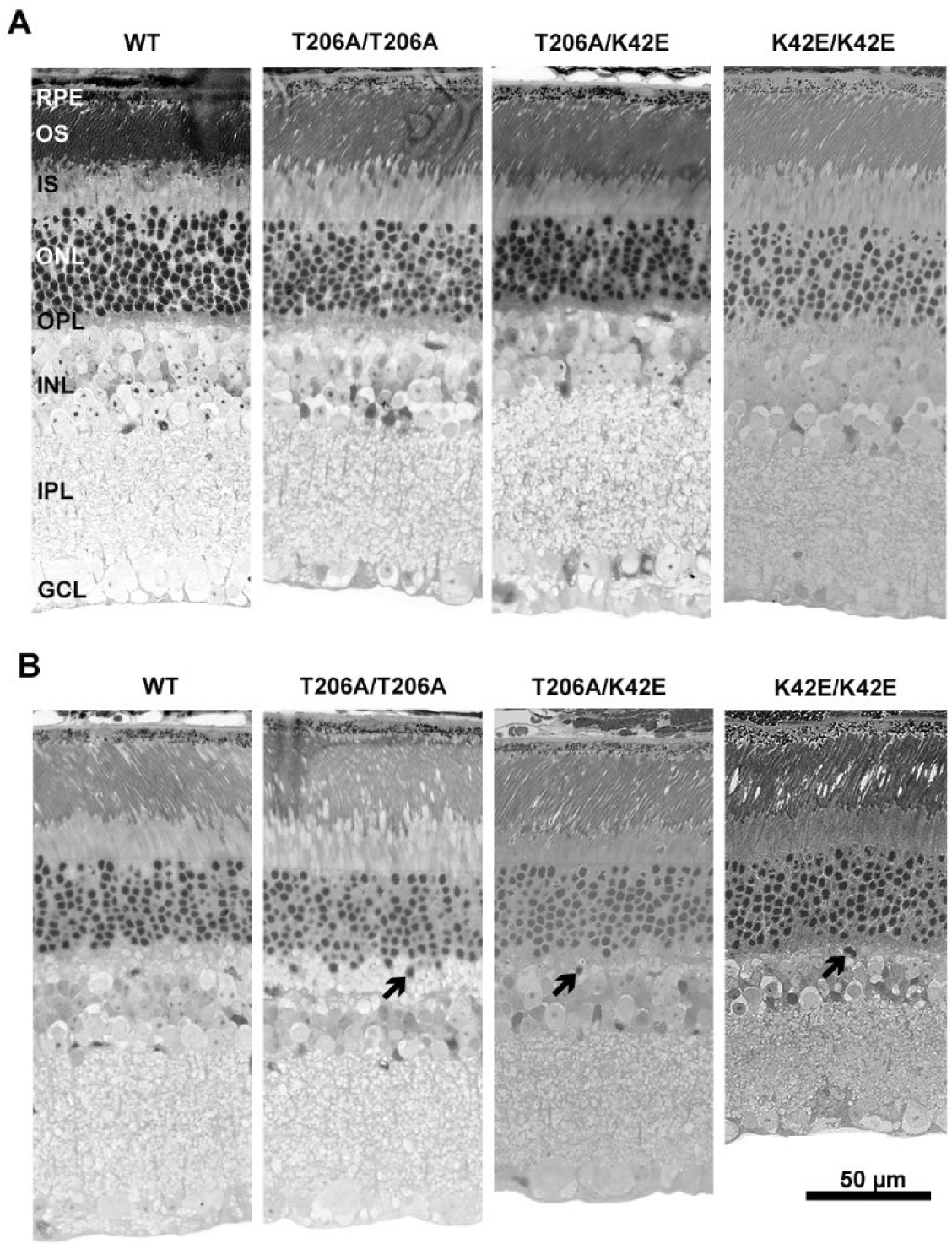
Retinal histology. Light micrographs of WT, T206A/T206A, T206A/K42E, and K42E/K42E mouse retina are shown for younger animals (PN ≤ 6-mo) in **(A)**; and for older animals (PN ≥ 12-mo) in **(B)**. INL thickness is reduced in all mutant strains, and ectopic migration of nuclei from the ONL into the OPL and INL is evident (*arrows*). *Abbreviations*: RPE, retinal pigment epithelium; OS, outer segment layer; IS, inner segment layer; ONL, outer nuclear layer; OPL, outer plexiform layer; INL, inner nuclear layer; IPL, inner plexiform layer; GCL; ganglion cell layer. Scale bar: 50 µm, all panels.

### The T206A *Dhdds* mutation causes thinning of the neural retina

To further characterize the effects of the T206A *Dhdds* mutation on retinal structure, we performed SD-OCT to quantify changes in total retinal thickness (TRT), and the thickness of the ONL and INL. TRT is measured from the internal limiting membrane (ILM, the vitreoretinal interface) and the boundary where the tips of the photoreceptor outer segments meet the apical face of the RPE. Representative SD-OCT images of WT, T206A/WT, T206A/T206A, and T206A/K42E mice are shown in **Fig. 3** at PN 1-mo (**Fig. 3A**) and PN 12-mo (**Fig. 3B**). Average TRT for WT, T206A/WT, T206A/T206A, and T206A/K42E mice were determined at PN 1-mo and PN 12-mo (**Fig. 3C**). There were no overt differences among the T206A/WT, T206A/T206A, nor T206A/K42E mice compared to their age-matched WT controls at either time point. However, there were statistically significant decreases in TRT at PN 12-mo compared to PN 1-mo: WT had a 5% (*p≤*0.001) reduction, T206A/WT had a 3% reduction (*p≤*0.05), T206A/T206A showed a 5% reduction (*p≤*0.01), and T206A/K42E had a 3% reduction (*p≤*0.05) in TRT. Previously reported TRT measurements for K42E/K42E are included for comparison (**Fig. 3C**) (Nguyen et al., 2023).

**Figure 3.**
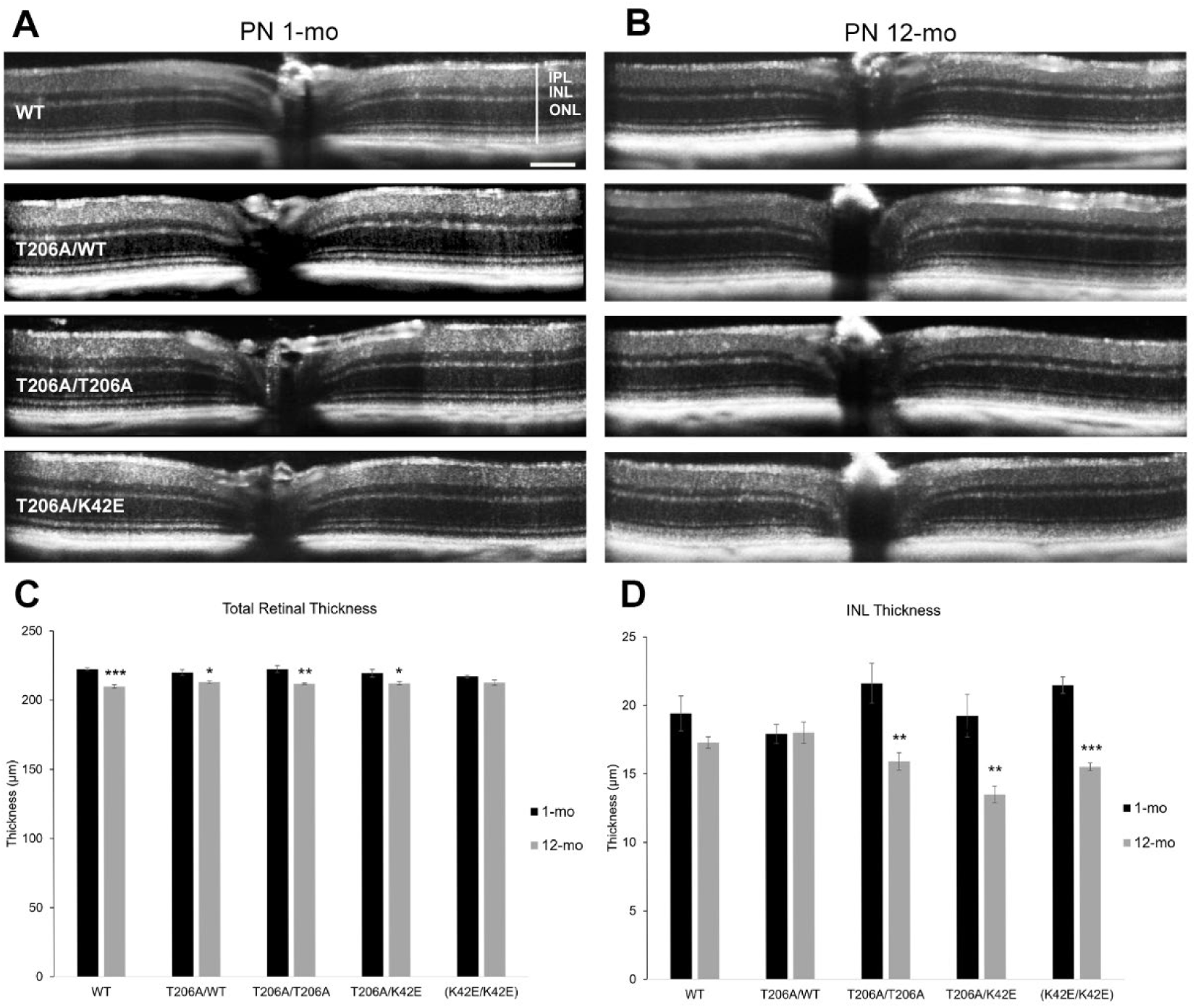
Measurements of total retinal thickness (TRT) and INL thickness obtained with SD-OCT. Representative images of WT, T206A/WT, T206A/T206A, and T206A/K42E SD-OCT scans at **(A)** PN 1-mo and **(B)** PN 12-mo. Total retinal thickness **(C**, measured by representative vertical white line in WT image in **(A))** and INL thickness **(D)** were averaged, graphed, and compared among PN 1-mo (*black*) and PN 12-mo (*grey*) mice. Previously published data obtained from K42E/K42E mice shown in panels C and D are included for reference (Nguyen et al., 2023). *Abbreviations* (see WT image in panel **(A))**: IPL, inner plexiform; INL, inner nuclear layer; ONL, outer nuclear layer. Scale bar: 0.1 µm for all panels. Statistical significance: \**p≤*0.05, \*\**p≤*0.01, and \*\*\**p≤*0.0001. WT N=4; T206A/WT, T206A/T206A, and T206A/K42E N=3.

Measurements of INL thickness were compared and graphed in **Fig. 3D**. At PN 1-mo, none of the mutant mouse strains showed any significant difference in INL thickness compared to WT. Thickness of the INL did not differ between PN 1-mo and PN 12-mo for either WT or T206A/WT mice. In contrast, T206A/K42E mice showed a significant 22% reduction in INL thickness at PN 12-mo compared to its WT counterpart (*p≤*0.001), and a significant 30% reduction compared to its PN 1-mo counterpart. Previously published K42E/K42E INL data showing significant reduction in INL thickness included for reference in **Fig. 3D** (Nguyen et al., 2023). Comparison of ONL thickness within the PN 1-mo and PN 12-mo time points and across the 12-mo time frame showed no significant differences (data not shown).

### The T206A mutation in *Dhdds* impairs photoreceptor-to-bipolar cell synaptic transmission

Representative PN 6-mo ERG waveforms for WT, T206A/WT, T206A/T206A, T206A/K42E mice are shown in **Fig. 4A** (dark-adapted, DA) and **Fig. 4B** (light-adapted, LA). The b-wave-to-a-wave (b/a) amplitude ratios were averaged and plotted for DA (**Fig. 4C**) and LA (**Fig. 4D**) conditions. Under DA (rod-driven) conditions, T206A/WT and T206A/T206A mice showed no difference compared to WT at PN 1-mo. However, the b/a ratio for T206A/WT mice was significantly higher than WT at PN 3- and 6-mo (*p≤*0.05). In contrast, T206A/K42E mice showed a significant decrease in b/a ratio (1.6 ± 0.03) compared to WT (1.9 ± 0.1) under DA conditions at PN 1-mo (*p≤*0.01). From PN 3-mo to PN 12-mo, the b/a ratios for T206A/T206A and T206A/K42E mice were progressively reduced compared to age-matched WT counterparts (*p≤*0.001 for both genotypes at all time points). The T206A/K42E b/a ratio was at the “negative ERG” threshold of b/a ratio =1 (Tanimoto et al., 2015) at PN 12-mo (1.0 ± 0.1, *p≤*0.001).

**Figure 4.**
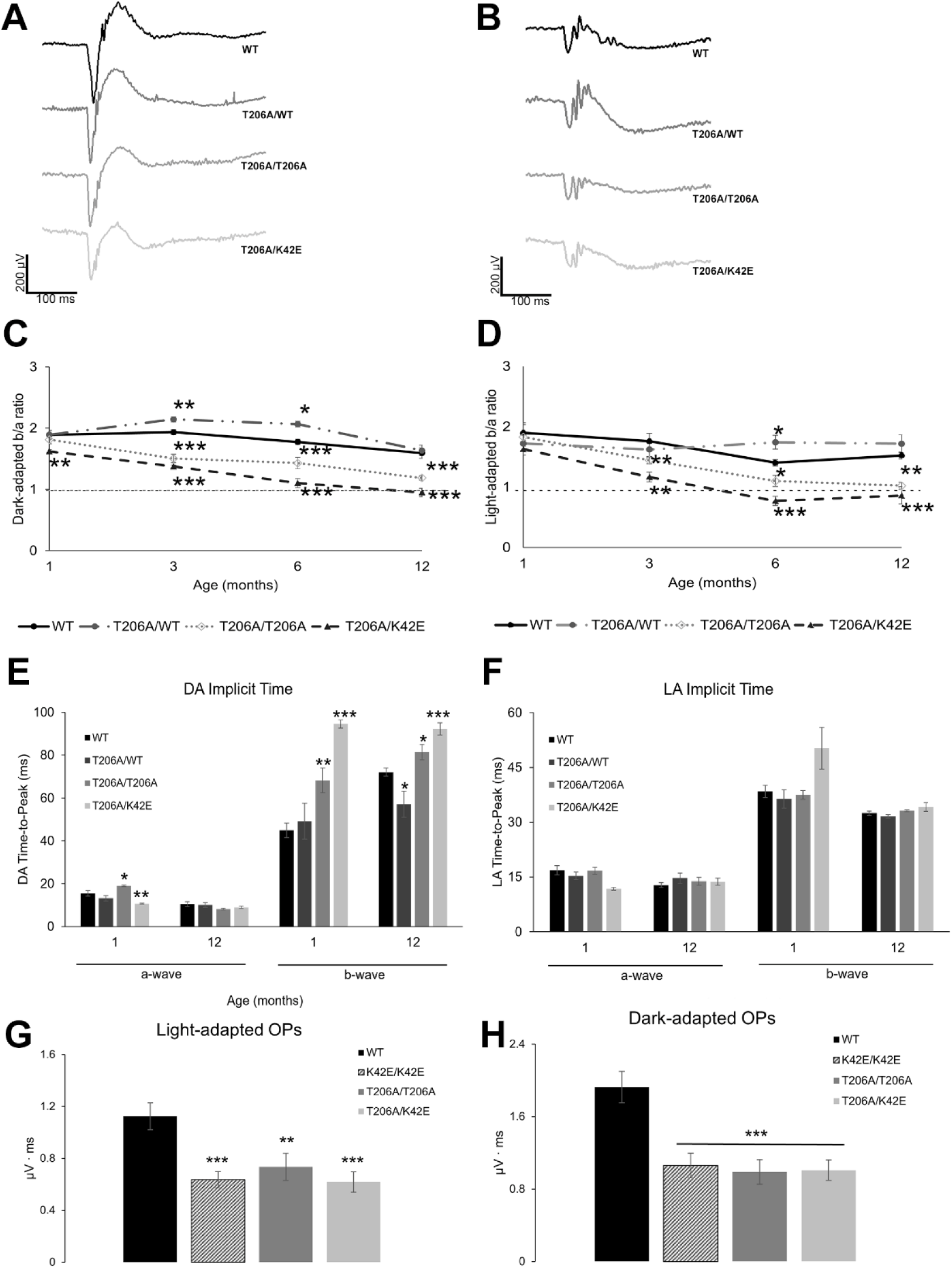
ERG b/a-wave amplitude ratios. Representative PN 6-mo **(A)** DA waveforms and **(B)** LA waveforms. The b/a ratios were taken at PN 1-, 3-, 6-, and 12-mo time points under **(C)** DA and **(D)** LA conditions. All mutant mouse b/a ratio comparisons were made in relation to WT values within each time point. WT (*solid black line*); T206A/WT (*dashed/dotted grey line*); T206A/T206A (*dotted grey line*); T206A/K42E (*dashed grey line*). The horizontal line at b/a=1 indicates the “negative ERG” threshold. **(E)** DA and **(F)** LA implicit time (time-to-peak) measurements of a-waves and b-waves of mutants are compared to WT at PN 1- and PN 12-mo. **(G)** DA and **(H)** LA oscillatory potential measurements are compared to WT at PN 12-mo. Statistical significance: *p<0.05, **p<0.01, and \*\*\**p≤*0.001. Animal numbers varied from N=6-14.

Under LA (cone-driven) conditions, the T206A/T206A and T206A/K42E b/a ratios did not differ from WT until PN 3-mo (*p≤*0.01). At PN 6-mo, however, the T206A/K42E b/a ratio dropped below the “negative ERG” threshold of b/a=1 and stayed below that threshold at PN 12-mo. Unlike the K42E/K42E mice (Nguyen et al., 2023), whose ERG response amplitudes reached “negative ERG” levels around PN 8-mo (DA) and PN 2-mo (LA), response amplitudes of T206A/T206A mice did not reach “negative ERG” levels under either DA or LA conditions up to PN 12-mo. However, the compound heterozygous T206A/K42E mice met the “negative ERG” criterion under DA (1.0 ± 0.1) and LA conditions (0.8 ± 0.1) at PN 12-mo. Raw ERG a-wave, b-wave, and b/a ratio values are listed in **Suppl. Table 1**.

Implicit time, or time-to-peak (TTP) amplitude after stimulus presentation, was measured for a-waves and b-waves under DA (**Fig. 4E**) and LA (**Fig. 4F**) conditions. Under DA conditions, the a-wave TTP for T206A/T206A and T206A/K42E mice differed significantly from WT at PN 1-mo but, surprisingly, did not differ at PN 12-mo. As previously reported for K42E/K42E animals (Nguyen et al., 2023), DA b-wave TTP for the mutant animals was significantly higher compared to WT mice at both PN 1- and 12-mo. However, under LA conditions, implicit times did not differ significantly from WT at either timepoint.

Oscillatory potentials (OP), or small wavelets on the rising phase of the b-wave, were analyzed by calculating the area under the curves. Averaged OPs are graphed for DA (**Fig. 4G**) and LA (**Fig. 4H**) conditions. All *Dhdds* mutants had significantly lower OP amplitudes (at least 40% reduction) compared to WT.

### Reduced c- and d-wave amplitudes indicate inner retinal dysfunction

Amplitudes of the ERG c-wave, which represent RPE and Müller cell contributions (Steinberg, 1985; Zeumer C, 1994) to the overall ERG response, were analyzed from PN 1-mo to PN 12-mo. Representative PN 6-mo waveforms are shown in **Suppl. Fig. 1A**. The c-wave amplitudes were averaged and graphed to compare each mutant mouse line against WT for each time point (**Suppl. Fig. 1B**). At PN 1-mo, c-wave amplitudes did not differ among the genotypes. At PN 3-mo, c-wave amplitudes for T206A/T206A and T206A/K42E mice were significantly lower than for WT mice (*p≤*0.05). In contrast, the c-wave amplitudes for T206A/WT mice were not significantly different from those of WT mice. At both PN 6- and 12-mo, c-wave amplitudes for all mutant genotypes were significantly lower than those of WT (*p≤*0.05).

ERG d-wave amplitudes (see *arrowheads*, **Suppl. Fig. 1A**) were measured and analyzed (**Suppl. Fig. 1C**) to assess the OFF-cone bipolar cell response (Naarendorp and Williams, 1999). Similar to c-wave analysis at PN 1-mo, d-wave amplitudes for mice of all mutant genotypes showed no differences compared to age-matched WT mice. However, from PN 3-mo onward, T206A/K42E mice had reduced d-wave amplitudes compared to age-matched WT mice (46.8% reduction, *p≤*0.05). By PN 6-mo, d-wave amplitudes of T206A/T206A mice also were significantly reduced (*p≤*0.05) compared to WT, and by PN 12-mo, mice of all mutant genotypes showed reduced d-wave amplitudes compared to WT.

### All *Dhdds* mutants exhibit significant reduction of bipolar and amacrine cell populations

Because all mutant mice retinas analyzed by ERG, histology and SD-OCT showed alterations consistent with inner retina compromise, we further explored structural changes in the inner retina of ≥PN 8-mo mutant mice (**Fig. 5**). Triple immunolabeling with bipolar cell type specific antibodies (CHX10, all bipolar cells (BPs); Islet-1, ON BPs (ISL1); PKC-α, rod BPs (PKC-α); also see table 1) was used to assess the integrity of these distinct bipolar cell populations. For analysis, BPs were classified by immunolabeling signature in each *Dhdds* mutant line and compared to wildtype controls to assess the effect of the different *Dhdds* mutations on rod, ON-cone, and OFF-cone BP classes (**Fig. 5Q**).

**Figure 5.**
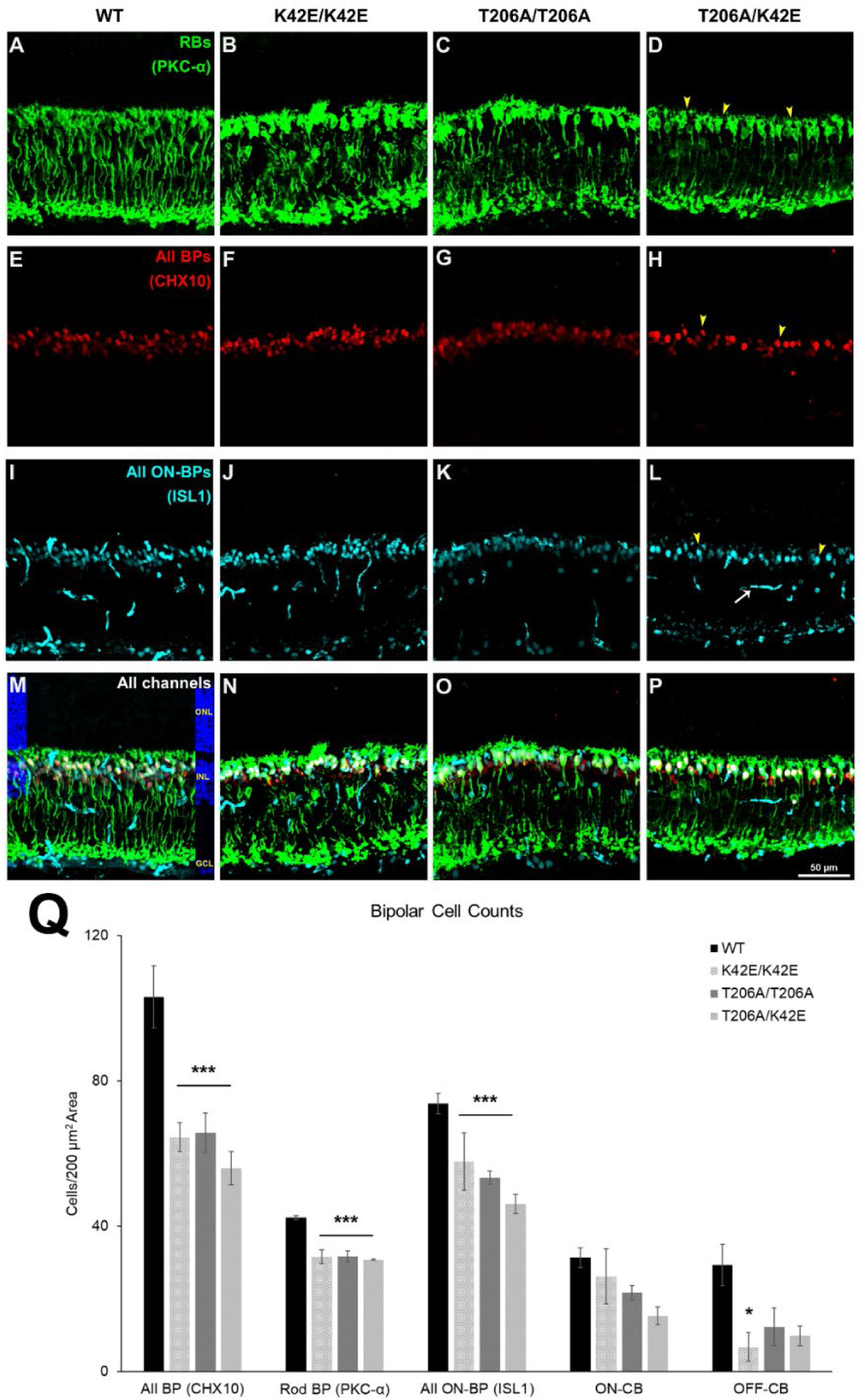
Bipolar cell densities. Representative 200 µm x 200 µm squares from retinal tissue sections are shown as obtained from WT (column 1), K42E/K42E (column 2), T206A/T206A (column 3), and T206A/K42E (column 4). Rod bipolar cells (RBs), labeled with anti-PKC-α, are shown in *green*, panels **(A)**-**(D)**; all BPs, labeled with anti-CHX10, are shown in *red*, panels **(E)**-**(H)**; all ON-BPs, probed with anti-ISL1, are shown in *cyan*, panels **(I)**-**(L)**; and merged channels with DAPI overlay **(M)** are shown in panels **(M)**-**(P)**. *Yellow arrowheads* in panels **(D)**, **(H)**, and **(L)** represent puncta that were counted. *White arrow* in panel **(L)** indicates ISL1-positive blood vessels (excluded from bipolar cell counts). Averaged overall values for rod BPs, ON-BP, ON-CB, and OFF-CB counts per 200 µm x 200 µm squares are graphed in panel **(Q)**. *Abbreviations*: As in legend, Figure 2. Scale bar: 50 µm for all panels. Statistical significance: \**p≤*0.05 and \*\*\**p≤*0.001. WT, K42E/K42E, T206A/T206A, and T206A/K42E N=3.

**Table 1.** Antibodies.

| Antibody | Host | Dilution | Vendor, Cat No. | RRID |
| --- | --- | --- | --- | --- |
| Chx-10 | Sheep,<br>polyclonal | 1:200 | GeneTex, GTX16142 | AB_368709 |
| Islet-1 | Mouse,<br>monoclonal | 1:100 | Developmental Studies<br>Hybridoma Bank,<br>39.4D5 | AB_2314683 |
| PKC-α | Rabbit,<br>polyclonal | 1:200 | Abclonal, A0267 | AB_2757080 |
| MEIS2 | Mouse,<br>monoclonal | Undiluted | Developmental Studies<br>Hybridoma Bank, 2B4 | AB_2618844 |
| TCF4 | Rabbit,<br>polyclonal | 1:150 | Proteintech, 22337-1-AP | AB_2879076 |
| Donkey anti-Sheep<br>Alexa Fluor 647 | Donkey/<br>IgG | 1:500 | Invitrogen, A21448 | AB_2535865 |
| Donkey anti-Mouse<br>Alexa Fluor 488 | Donkey/<br>IgG | 1:500 | Invitrogen, A21202 | AB_141607 |
| Donkey anti-Rabbit<br>Alexa Fluor 546 | Donkey/<br>IgG | 1:500 | Invitrogen, A10040 | AB_2534016 |

Analysis of total CHX10+ labeling showed significant BP loss in all three *Dhdds* mutant strains compared to WT mice (K42E/K42E, 36% reduction; T206A/T206A, 36% reduction; T206A/K42E, 43% reduction, *p≤*0.001 for all comparisons, **Fig. 5Q**). Analysis of PKC immunolabeling revealed significant rod bipolar cell loss in all *Dhdds* mutant lines compared to WT mice (K42E/K42E, 29% reduction; T206A/T206A, 23% reduction; T206A/K42E, 27% reduction. *p≤*0.001 for all comparisons, **Fig. 5Q**).

To analyze the effects of the various *Dhdds* mutations on the ON-cone BPs population, we subtracted the numbers of PKC+ rod BPs from the total number of Islet-1+ ON BPs in each specimen to determine the numbers of ON-cone BPs for each mouse strain. Reduced numbers of ON-cone BPs were noted in each *Dhdds* mutant strain although the difference was not statistically significant from WT mice: K42E/K42E, 16% reduction; T206A/T206A, 31% reduction; T206A/K42E, 51% reduction compared to WT (**Fig. 5Q**). To determine whether *Dhdds* mutations affected the OFF-cone BP population, the total number of ISL1+ bipolar cells (rod BPs + ON-cone BPs) was subtracted from the total number of BPs (Chx10+ cells) in each specimen from each *Dhdds* mutant mouse strain and compared to WT controls: K42E/K42E, 77% reduction; T206A/T206A, 58% reduction; T206A/K42E, 66% reduction. The reduction in OFF-cone BPs differed significantly from WT only in the K42E/K42E retina (*p≤*0.05).

Given the thinning of the INL and reduced bipolar cell populations in *Dhdds* mutants, we also examined the amacrine cell (AC) population. To identify ACs, we used MEIS2, a transcription factor for GABAergic ACs, and TCF4, a transcription factor for glycinergic ACs (Yan et al., 2020). Representative double immunofluorescence images of labeling for MEIS2 and TCF4 are shown in **Fig. 6A-D**, with MEIS2 pseudo-colored red, and TCF4 pseudo-colored green. Cells were counted in the same manner as BPs. Averaged GABAergic and glycinergic AC populations for each genotype are graphed in **Fig. 6E**. All *Dhdds* mutant mice had reduced numbers of GABAergic and glycinergic ACs compared to age-matched WT mice. T206A/K42E mice had the largest difference compared to WT, with a 25% AC loss (*p*≤0.001) in both AC types; K42E/K42E showed 20% AC loss (*p*≤0.001). In comparison, the T206A/T206A mutants showed less severe AC loss, with 10% dropout of glycinergic AC and 14% loss of GABAergic AC (*p*≤0.001, both).

**Figure 6.**
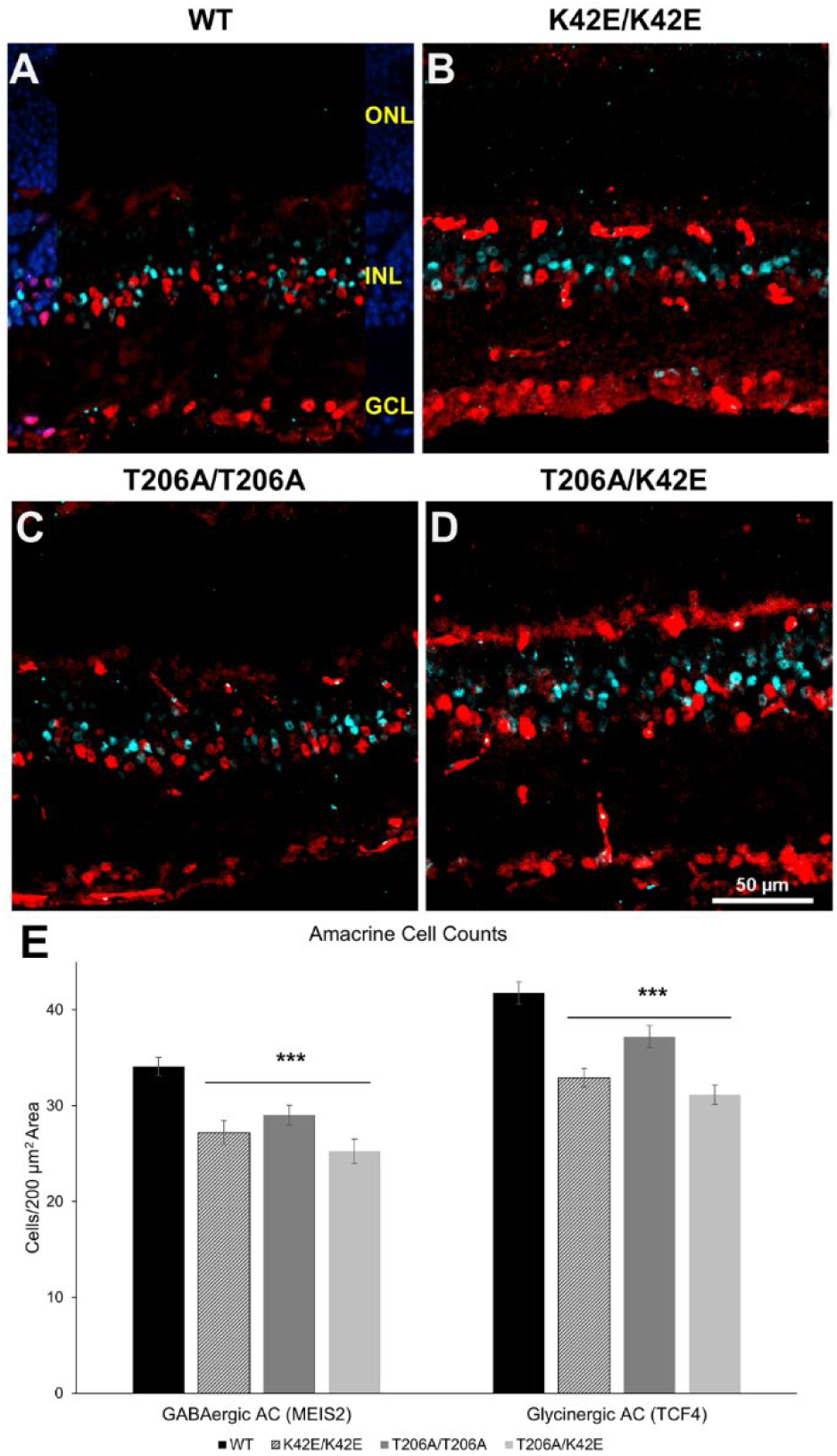
Amacrine cell densities. Representative 200 µm x 200 µm squares from immunolabeled retinal tissue sections are shown from WT **(A)**, K42E/K42E **(B)**, T206A/T206A **(C)**, and T206A/K42E **(D)** mice. GABAergic AC nuclei show MEIS2 labeling (pseudo colored red) and glycinergic AC nuclei show TCF4 labeling (pseudo colored cyan). A DAPI panel overlay is shown to illustrate the nuclear layers of the retina **(A)**. Averaged overall values for GABAergic and glycinergic AC nuclei counts are graphed in panel **(E)**. *Abbreviations*: same as legend, Figure 2. Scale bar: 50 µm for all panels. Statistical significance: \**p≤*0.05 and \*\*\**p≤*0.001. WT, K42E/K42E, T206A/T206A, and T206A/K42E, N=3.

### Visual acuity is greatly reduced in *Dhdds* mutant mice relative to WT controls

The optokinetic response (OKR) was examined at PN 12-mo to assess the status of signal transmission from the retina to the brain. Spatial frequency was measured, the highest values were recorded, and visual acuity values were graphed under LA conditions (**Suppl. Fig. 2A**) and under DA conditions (**Suppl. Fig. 2B**). Compared to WT controls, T206A/T206A and T206A/K42E mice both showed significantly lower visual acuity under both LA and DA conditions (*p≤*0.05).

## DISCUSSION

We generated and characterized a novel mouse line that harbors one of the disease-associated *DHDDS* mutations found in RP59 patients. Our goal was to create a tractable and informative RP59 animal model to better understand the underlying mechanism leading to the human disease. Previously, we studied the K42E/K42E knock-in mouse model, based upon the primary *DHDDS* mutation found in RP59 patients, where we found reduced total retinal and INL thicknesses, photoreceptor terminal retraction, second order neuronal "sprouting," ectopic cell displacement, and “negative ERG” waveforms without any evidence of defective protein N-glycosylation (Nguyen et al., 2023; Ramachandra Rao et al., 2020). In the present study, relevant to the second most prominent *DHDDS* mutation found in RP59 patients, T206A, we also assessed the structural and functional effects of the mutation on the retina. As we hypothesized, the T206A/K42E knock-in mice expressed comparable phenotypes to those of K42E/K42E knock-in mice. Moreover, independent from the K42E allele, mice with the T206A/T206A mutation (not reported found in RP59 patients), also exhibit a similar phenotype. Knock-in mice showed declining structural and functional phenotypes with age, but most importantly, also showed reduced bipolar cell densities, indicating increased bipolar cell sensitivity to insult as a result of DHDDS mutation.

As expected with retinal aging, total retinal thickness was reduced at PN 12-mo, but the INL thickness in the T206A/T206A and T206A/K42E mice showed significant reductions by 26% and 30%, respectively (compared to 11% in age-matched WT mice, **Fig. 3**). Histological analysis of retinal tissue sections performed on younger animals (PN ≤ 6-mo) showed no compromise of gross retinal structure in T206A/T206A or T206A/K42E mice. However, older T206A/T206A and T206A/K42E mice (PN ≥ 6-mo) exhibited reduced total retinal thickness (TRT) as well as displacement of photoreceptor cells from the ONL (*arrows*, **Fig. 2**). Previously published data from K42E/K42E mice (Nguyen et al., 2023; Ramachandra Rao et al., 2020) showed increased GFAP immunolabeling, increased cell death (TUNEL staining), and photoreceptor terminal retraction and "sprouting" of rod bipolar cell dendrites into the ONL (Nguyen et al., 2023; Ramachandra Rao et al., 2020). However, there were no significant changes in ONL thickness, indicating that the aforementioned histological changes were not merely due to increasing age and were restricted to the *inner* retina, rather than the photoreceptor layer. The T206A allele has a milder degenerative phenotype than the K42E allele, and in mice, both alleles are less severe than that reported for RP59 patients (Kimchi et al., 2018). Many phenotypes found in patients pointed to issues in and around the fovea, which cannot be addressed in our study, since mice lack a macula or fovea.

Are the RP59 knock-in mice good models for the human disease? The primary difference between the disease phenotypes observed is that mice show degenerative changes in the inner retina (bipolar and amacrine cells) and the primary site of cell loss in patients is in the outer segments (rod photoreceptors). As shown in **Table 2**, the genotypes have several common attributes. Also note that the average best corrected visual acuity for RP59 patients examined to date ranges from 20/20 to 20/400 (Fliesler et al, 2022). Thus, there is significant useful vision remaining in most of the patients. It is possible that the same mechanism of disease is operating in both patients and animal models, but the phenotypic outcome is different due to species differences (largely the presence or absence of a macula).

**Table 2.**
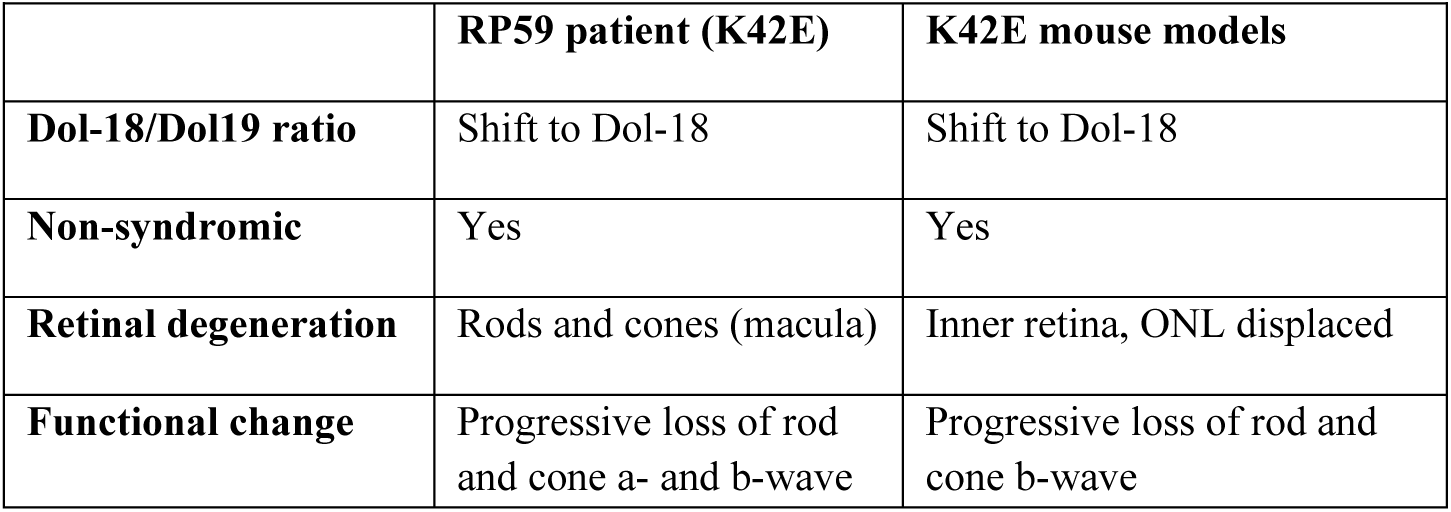
Comparison of RP59 patient and mouse model phenotypes.

| Table 2. Comparison of RP59 patient and mouse model phenotypes. |  |  |
| --- | --- | --- |
|  | RP59 patient (K42E) | K42E mouse models |
| Dol-18/Dol19 ratio | Shift to Dol-18 | Shift to Dol-18 |
| Non-syndromic | Yes | Yes |
| Retinal degeneration | Rods and cones (macula) | Inner retina, ONL displaced |
| Functional change | Progressive loss of rod and cone a- and b-wave | Progressive loss of rod and cone b-wave |

Because of the histological thinning of the INL and evidence of inefficient synaptic transmission to second-order neurons evidenced by ERG, we assessed BPs using immunofluorescence imaging. The cell bodies of BPs make up 40-50% of the INL (Wassle et al., 2009). Upon visual inspection, it was apparent that there were fewer numbers of CHX10+ cells in mutants compared to WT (**Fig. 5E-H**). Overall, all mutants had fewer BPs per 200 µm x 200 µm square, with T206A/K42E mice showing the biggest loss at 46% of the WT numbers. The most fascinating piece of data we found from BP density analysis was the significant (and selective) reduction of ON BPs (depicted by PKC-α+ and ISL+ cells, **Fig. 5**), which correlates directly to the observed “negative ERG” and decreased sensitivity (reflected by increased implicit times, **Fig. 4**). The reduced b-wave amplitudes and decreased sensitivity under dark-adapted conditions (**Fig. 4C and E**) correlate directly to the significant reduction of rod BP densities, as the b-wave is initiated from ON BPs (Bhatt et al., 2023; Creel, 2019). While mutant rod BP counts were reduced compared to those of WT, ON-cone BP counts were not. This is consistent with the observed reduction of scotopic b-wave amplitudes with no concomitant difference in sensitivity under light-adapted conditions (**Fig. 4F**). Although we did not probe directly for OFF-cone BPs, we calculated estimates by taking the numbers of ISL1+ cells and subtracted from the total number of CHX10+ cells, which revealed a marked and significant reduction in K42E/K42E OFF cone BPs (77% fewer than WT, **Fig. 5Q**). However, T206A/T206A and T206A/K42E OFF-cone BP counts were not statistically lower than WT. Reduced OFF-cone BP counts are in agreement with the reduced d-wave ERG amplitudes found in mutant mice (**Suppl. Fig. 1**) (Perlman, 1995). Currently, we do not have a precise mechanism to explain why there is a drop-out of bipolar cells.

Because the inner retina is very thin in older *Dhdds* mutant mice and the reduction in thickness corresponds to the region of the INL occupied by ACs (see **Fig. 2** and **3**), loss of these cells was expected (**Fig. 6**). It is estimated that about 50% of ACs in mammalian retinas are GABAergic, about 40% of ACs are glycinergic, and are critical to processing of signals transmitted to ganglion cells (Davanger et al., 1991; Perez-Leon et al., 2022). Therefore, labeling for MEIS2 (GABAergic ACs) and TCF4 (glycinergic ACs) together should identify roughly 90% of the total AC population. All *Dhdds* mutant strains showed reduced AC density, consistent with a reduction in BP densities. Further, these data are congruent with the reduced oscillatory potentials (**Fig. 4G-H**), which arise from inhibitory pathways mediated by GABAergic and glycinergic ACs in the IPL (Wachtmeister, 1998). Thus, the reduced oscillatory potentials observed in the current studies suggest that loss of GABAergic and glycinergic ACs in *Dhdds* mutant mice significantly impairs synaptic processing by amacrine cells in the IPL.

To test whether T206A *DHDDS* mutations caused dysfunction in transmission of light-driven visual signals to the brain, we performed OKR experiments on PN 12-mo mice (**Suppl. Fig. 2**). Unlike humans, mice have laterally positioned eyes and lack the ability to rotate their eyes. OKR (which is performed in awake and unrestrained animals, allowing head movement) provides an accurate, quantitative method to assess visual competence, as represented by calculation of numerical values for visual acuity from spatial frequency measurements, or ability to resolve two points in space (Stahl, 2004). We found that both the T206A/T206A and T206A/K42E knock-in mouse lines have reduced visual acuity (*i.e.*, greater spatial frequency values), compared to age-matched WT mice. Compared to previously published OKR data (Nguyen et al., 2023), the K42E/K42E mice had lower light-adapted visual acuity than the T206A/K42E and T206A/T206A mice. It is likely that the loss of bipolar cells and impaired transmission of signals through the inner retina arising from the K42E and T206A mutations together impair the visual performance of our *Dhdds* mutant mice.

As shown previously, K42E/K42E *Dhdds* mutant mice display a negative ERG as early as PN 2-mo (Nguyen et al., 2023). In RP59 patients, the T206A *Dhdds* mutation only has been found, to date, in heterozygous combination with the K42E mutation (Wen et al., 2013). The K42E/K42E *Dhdds* mutation caused a severe phenotype in the knock-in mice, while the T206A/K42E knock-in mutants exhibited a similar, but less severe, phenotype. Apparently, the K42 residue, which is in direct contact with the CPT active site, is particularly important for the activity of the protein (Bar-El et al., 2020; Lisnyansky Bar-El et al., 2019). As found in human patients and in the K42E/K42E mouse model, the K42E mutation led to shortened dolichol species (Nguyen et al., 2023; Wen et al., 2013). The hypothesis that the positively charged K-to-E amino acid mutation hinders the active site of the CPT enzyme complex is supported by the more severe phenotype in the K42E/K42E mice, followed by a less severe phenotype in our T206A/K42E mice (Lisnyansky Bar-El et al., 2019). A milder phenotype is shown in the T206A/T206A mice. The T206 residue is also hypothesized to perturb the active site, as its threonine backbone is hydrogen-bonded with the backbone of the metal-binding D34 residue (Edani et al., 2020). However, the question of why the mutations lead to non-syndromic retinitis pigmentosa remains unresolved.

In summary, structural and functional compromise of the inner retina in RP59 mouse models are the primary phenotypic features observed, consistent with defective photoreceptor to inner retinal neuronal transmission. While the precise mechanism that leads to synaptic dysfunction remains to be determined, we speculate that gain of function in cPT (causing formation of dolichol species having shorter than normal chain lengths) leads to loss of function of one or more synaptic proteins. This could occur through mislocalization of DHDDS (or cPT holoenzyme) to the synapse and subsequent ectopic interaction with synaptic protein(s) or, perhaps more likely, through the ectopic interaction of DHDDS/cPT with synaptic protein(s) in the ER or Golgi where all of these proteins are normally found to facilitate postranslational modification. Direct proteomic analysis and high-resolution immunocytochemistry will be needed to confirm or refute this speculation and to more definitively elucidate the precise disease mechanism.

## MATERIALS AND METHODS

### Animals

*Dhdds*^T206A/WT^ mice were created on a C57Bl/6J background by the UAB Heflin Genomics Center using CRISPR-Cas9 technology. Briefly, CRISPR guides were designed using CRISPOR (http://crispor.tefor.net) to target exon 7 in the mouse *Dhdds* locus using the following amplicon with a silent DNA polymorphism to eliminate the PAM recognition site downstream of the target site for DNA cleavage (Anders et al., 2014). Founder animals were identified by PCR using primers flanking the target loci that amplified a 399 base pair fragment in wild type (WT) animals. Founder animals were identified by PCR using primers flanking the target loci that amplified the 399 base pair fragments in WT animals.

Sequence-verified heterozygous mice were crossed to create WT and homozygous *Dhdds*^T206A/T206A^ mice. Homozygous *Dhdds* T206A/T206A mice were crossed with homozygous *Dhdds* K42E/K42E mice to create compound heterozygous T206A/K42E mice. Genotype of cross-bred mice was confirmed by PCR and DNA sequencing (see *Methods-PCR Genotyping*). All procedures conformed to the *ARVO Statement for the Use of Animals in Ophthalmic and Vision Research* and were approved by the Institutional Animal Care and Use Committee (IACUC) of the University of Alabama at Birmingham.

Animals were maintained on a standard 12 h/12 h light/dark cycle (20–40 lux ambient room illumination), fed standard rodent chow, provided water *ad libitum*, and were housed in plastic cages with standard rodent bedding.

### PCR Genotyping

Initial genotype verification was completed at the UAB Heflin Center for Genomic Sciences Core Laboratories, validating the heterozygous T206A/WT strain. PCR primers were purchased from Invitrogen (Waltham, MA, USA) to validate tail snip DNA sequences: (forward primer, 5’-TGGGTGAT-CTGCATCTGCTG-3’ and reverse primer, 5’-GTGCACCATGGTTCCTCTGA-3’). DNA sequences were confirmed in subsequent generations by restriction enzyme digestion with BclI (for T206A) and StyI (for K42E), which cleave the respective knock-in alleles.

WT amplicon (Thr206 in **bold**): TGGGTGATCTGCATCTGCTGCCCTTGGACCTCCAGGA-GAAGATTGCGCATGCCATCCAGGCTACTAAGAACTACAATAAGTGTTTCCTCAATGT CTGCTTTGCATACACATCACGTCATGAGATTGCCAATGCTGTGAGAGAGATGGCCTG GGGCGTGGAACAAGGTCTGCTGGAACCCAGTGATGTCTCCGAGTCTCTGCTCGATAA GTGCCTCTATAGCAACCACTCTCCTCATCCCGACATCCTGATCCGG**<u>ACT</u>**TCTGGGGAGGTGCGGCTGAGTGACTTCTTGCTCTGGCAGACGTCCCATTCCTGCCTCGTGTTCCAG CCTGTCCTGTGGCCAGAATACACATTTTGGAACCTGTGTGAGGCAATTCTGCAGTTTCAGAGGAACCATGGTGCAC; 200 bp ssDNA oligo repair template (A206 in **<u>bold</u>**; BclI restriction site unique to mutant allele in *italics*\*): CTGTGCTTCTGTCTCCTGCCCACCTAG-TGATGTCTCCGAGTCTCTGCTCGATAAGTGCCTCTATAGCAACCACTCTCCTCATCCC GACATCC*TGATCa*GG**<u>gCT</u>**TCTGGGGAGGTGCGGCTGAGTGACTTCTTGCTCTGGCAGG TAGGTTGTTTTGAAACATGTTATTTTGGGGTTGGGCTGAACCCTGGAACTGAAGCAG *CGG>AGG silent mutation at CRISPR PAM recognition site.

### Spectral domain optical coherence tomography (SD-OCT)

*In vivo* retinal imaging was performed using a Bioptigen Model 840 Envisu Class-R high-resolution SD-OCT instrument (Bioptigen/Leica, Inc), as described previously by DeRamus et al. (DeRamus et al., 2017). Retinal layer thickness values were measured using the Bioptigen (Leica, Inc.) Diver V 3.4.4 software and were spot checked manually using calipers on the Bioptigen (Leica, Inc.) InVivoVue software. Data were collected from at least N=3 independent WT, T206A/T206A, T206A/WT, and T206A/K42E mice at PN 1-mo and PN 12-mo.

### Optokinetic Response (OKR)

To assess how the brain is responding to visual stimuli, the optokinetic response (OKR) was measured as described in Gil et al. (Gil et al., 2022) under scotopic (dark-adapted) and photopic (light-adapted) conditions using an OptoMotry HD instrument (Cerebral Mechanics Inc.). Data were analyzed using Microsoft 365 Excel and visualized graphically.

### Electroretinography (ERG)

Mice were dark-adapted (DA) overnight and ERGs were recorded on a custom-built system as described previously (Nguyen et al., 2023). Briefly, ERG a- and b-wave responses were recorded following 2-ms flashes of light with increasing flash intensities under dark-adapted and light-adapted (LA) conditions, while ERG c- and d-waves were recorded following a 5-sec step of light stimulus using in DC recording mode. Implicit times for a- and b-wave responses were recorded at the brightest flash intensities.

Oscillatory potentials were analyzed for waves recorded following the highest light stimuli (6.955 log photons/µM^2^) under dark-adapted and light-adapted conditions. Methods were as described by Hancock and Kraft, 2004 (Hancock and Kraft, 2004). Briefly, a high pass filter of 70 Hz and a low pass filter of 34 Hz were applied to each wave. The area under the curve from time=0s to time =0.15s was calculated and noise was subtracted to quantify the amplitudes of oscillatory potentials.

Response amplitudes were averaged using Data Browser V 6.4.4 software (LabVIEW by National Instruments, Austin, TX, USA) and analyzed with IGOR Pro 8 & 9 (WaveMetrics, Inc., Portland, OR, USA).

### Light Microscopy (Histology)

Mouse eyes were enucleated following euthanasia and cervical dislocation, and then fixed in buffered mixed aldehydes, and processed for Epon 812 resin embedment as described in detail previously (Sarfare et al., 2014). Retinal tissue sections (0.8-µm thickness) were obtained using a microtome, collected on glass microscope slides, and stained with 0.1% Toluidine blue. Histological images were collected using an Olympus VS-120 photomicroscope (BX61VS platform) running Olympus VS-ASW-2.9 software.

### Immunofluorescence histochemistry

Mouse eyes were processed to obtain retinal cryosections as previously described (Wang et al., 2017). In brief, after euthanasia and enucleation, eyes were “fixed” by immersion in chilled 4% paraformaldehyde (PFA) in 0.1M PBS for 15 min, and the anterior segments were removed. The resulting eyecups were cryoprotected by serial immersion in graded buffered sucrose solutions, then embedded and frozen in O.C.T. (Optimal Cutting Temperature) compound. Cryosections (10-12 µm thickness) were obtained using a cryotome and were collected on glass microscope slides, then were rehydrated in PBS and “blocked” using 10% donkey serum in PBS containing 0.1% Triton-X 100 (PBST) for 1 h, before being incubated in primary and appropriate fluor-tagged secondary antibodies (see **Table 1** for antibody information). Following DAPI counterstaining, the slides were cover slipped. Eyes from N=3 different animals of each genotype (WT, *Dhdds*^K42E/K42E^, *Dhdds*^T206A/T206A^, and *Dhdds*^T206A/K42E^) between the age range of PN 8-mo to PN 12-mo were used. At least N=3 retinal sections (including or proximal to the optic nerve head) were subjected to immunofluorescence microscopy analysis, per genotype and age, and at least N=5 images (200 µm x 200 µm square) acquired from both sides of the optic nerve head were analyzed from each retinal section.

Labeled retinal cryosections were imaged using a Nikon AX-R Confocal Microscope at 20X magnification. Z-stack images were compressed, “denoised”, and analyzed using the NIS-Elements AR imaging software (Nikon Instruments, version 5.21.03). Labeled cell bodies within each 200 µm x 200 µm square were counted manually in each fluorescence channel and catalogued in the NIS-Elements Object Count tool. Cell counts were averaged and analyzed using R software (version 4.2.1).

### Statistical Analysis

Statistical analysis was done using a two-tailed Student’s *t*-test assuming equal variances or, alternatively, using ANOVA with *post-hoc* Tukey’s test to compare quantitative data obtained from T206A/T206A, T206A/WT, and T206A/K42E mice in comparison with age-matched WT mice. Outlier values were identified and removed using the interquartile range method. Statistical significance *p*-value thresholds were: \**p*<0.05, \*\**p*<0.01, and \*\*\**p*≤0.001.

## ACKNOWLEDGEMENTS

We thank Nicole Naylor for assistance with OKR; Isaac P. Cobb for technical assistance; Drs. Laura Lambert and Robert Kesterson of the UAB Transgenic & Genetically Engineered Models Core for generation of the T206A mice; Dr. Michael Crowley in the Heflin Center for Genomic Science Core Laboratories for DNA sequencing; Dr. Timothy Kraft and James Fortenberry in the Ocular Phenotyping Core for assistance with ERG; Gillian Huskin and Dr. Yuchen Wang for providing an immunofluorescence protocol and aliquot of secondary antibodies; Dr. Jose Luis Roig Lopez in the Molecular & Cellular Analysis Core for assistance with confocal microscopy; and Dr. Sriganesh Ramachandra Rao for helpful discussions throughout the course of this research.

## Funding

The research described herein by the authors was supported by U.S. Department of Health and Human Services (National Institutes of Health (NIH)/National Eye Institute (NEI)) grant R01 EY029341 to S.J.P. and S.J.F., and NIH/NEI core grant P30 EY003039 to S.J.P.; NIH/NEI F31

EY032764 fellowship (M.N.N); as well as support from the UAB Vision Science Research Center (S.J.P., M.N.N., D.C.), and facilities and resources provided by the VA Western NY Healthcare System (S.J.F.). S.J.F. is the recipient of a BLR&D Research Career Scientist Award (I K6 BX005787) from the Department of Veterans Affairs. Research reported in this manuscript also was supported, in part, by the National Center for Advancing Translational Sciences (NCATS) of the National Institutes of Health under award number UL1TR003096 (awarded to the University at Buffalo). Additional support (to S.J.F.) was provided by funds from a Paul Kayser International Award in Retina Research from Retina Research Foundation. Services obtained from the UAB Transgenic & Genetically Engineered Model Systems Core Facility in this publication were supported by awards NIH P30 CA013148, P30 AR048311, P30 DK074038, P30 DK056336, and P60 DK079626. The opinions expressed herein are not those of the Department of Veterans Affairs nor of any other U.S. Government agency.

### Data availability

All relevant data can be found within the article and its supplementary information.

## Supplementary information

The online version contains Supplementary material.

