## Supplemental Figs 1 and 2 for "Modeling Retinitis Pigmentosa 59: *Dhdds* T206A and *Dhdds* K42E knock-in mutant mice are phenotypically similar"

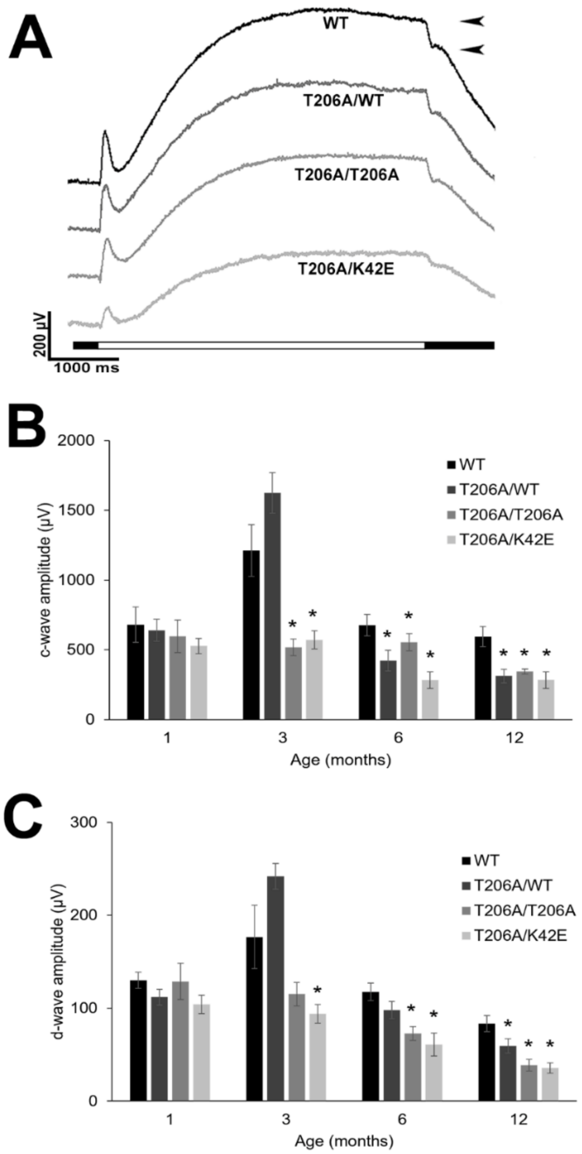


**Supplemental Figure 1. ERG c- and d-wave amplitudes. (A)** Representative PN 6-mo waveforms for c- and d-waves are shown with *black arrowheads* indicating measurement of d-wave amplitudes and horizontal bar indicating a 5000 ms stimulus on (*white bar*) and stimulus off (*black bar*). PN 1-, 3-, 6-, and 12-mo measurements for **(B)** c-wave amplitudes and **(C)** d-wave amplitudes and compared to WT values for each time point. Statistical significance: *$p \leq 0.05$, **$p \leq 0.01$, and ***$p \leq 0.001$. Animal numbers varied from N=6-14.

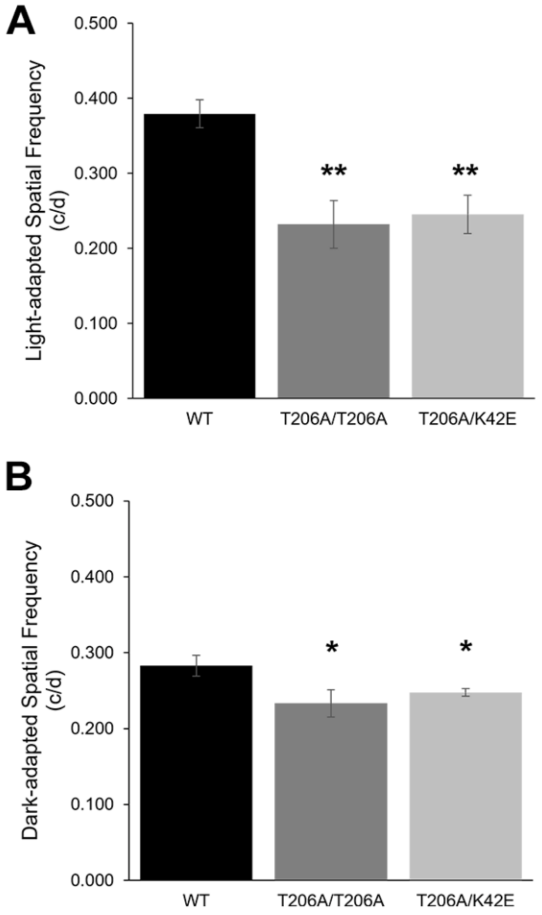


**Supplemental Figure 2. Visual acuity values at PN 12-mo.** Optokinetic responses (OKR) were evaluated to determine the highest spatial frequency and visual acuity was analyzed for WT, T206A/T206A, and T206A/K42E mice at PN 12-mo under **(A)** LA conditions and **(B)** DA conditions. T206A/WT visual acuities did not differ from WT (*not shown*). *Abbreviations*: c/d, cycles/degree. Statistical significance: *$p\leq0.05$ and **$p\leq0.01$. WT, T206A/T206A, and T206A/K42E $n\leq3$.
